# QOT: Efficient Computation of Sample Level Distance Matrix from Single-Cell Omics Data through Quantized Optimal Transport

**DOI:** 10.1101/2024.02.06.578032

**Authors:** Zexuan Wang, Qipeng Zhan, Shu Yang, Shizhuo Mu, Jiong Chen, Sumita Garai, Patryk Orzechowski, Joost Wagenaar, Li Shen

## Abstract

Single-cell technologies have emerged as a transformative technology enabling high-dimensional characterization of cell populations at an unprecedented scale. The data’s innate complexity and voluminous nature pose significant computational and analytical challenges, especially in comparative studies delineating cellular architectures across various biological conditions (i.e., generation of sample level distance matrices). Optimal Transport (OT) is a mathematical tool that captures the intrinsic structure of data geometrically and has been applied to many bioinformatics tasks. In this paper, we propose QOT (Quantized Optimal Transport), a new method enables efficient computation of sample level distance matrix from large-scale single-cell omics data through a quantization step. We apply our algorithm to real-world single-cell genomics and pathomics datasets, aiming to extrapolate cell-level insights to inform sample level categorizations. Our empirical study shows that QOT outperforms OT-based algorithms in terms of accuracy and robustness when obtaining a distance matrix at the sample level from high throughput single-cell measures. Moreover, the sample level distance matrix could be used in downstream analysis (i.e. uncover the trajectory of disease progression), highlighting its usage in biomedical informatics and data science.

## 1 Introduction

Single-cell technologies have enabled the opportunity to explore the molecular mechanism through high-dimensional gene analyses. Numerous Computational frameworks have recently been developed in order to vary between cells. For example, [1] developed PhenoGraph to study the single-cell heterogeneity. [2] developed SIMLR to improve the clustering performance of single-cell sequencing data as well as provide better visualization and interpretability.

Pathomics enhances molecular omics approaches by providing quantitatively measured, structure-focused details on histological formations, emphasizing objective, tissue-related data. [3] and [4] investigated extracting and analyzing computer-generated measurements from digitized images in histopathology.

Optimal transport (OT) is a mathematical concept initially proposed by [5] and subsequently reformulated in a more computationally-friendly manner by [6]. This concept has garnered considerable interest from theoretical and practical perspectives [7, 8]. In recent times, its applicability has been rediscovered and extended to fields such as image processing [9], shape analysis [10], generative modeling [11], and fairness [12]. At its core, OT addresses the challenges of comparing empirical distribution, ensuring mass preservation while minimizing associated costs. The application of Optimal Transport (OT) has recently been extended to similarity learning as a way to find the distance between two high-dimensional distributions [13, 14]. In particular, with the advancement of single-cell omics data such as RNA-seq and cytometry, OT has been used in trajectory identification, cell-cell similarity inference, and single-cell data auto-gating [15, 16, 17].

Until now, only a few methods have been characterizing the analysis of single-cell datasets at the sample levels. PhEMD (phenotypic earth mover’s distance) [18] used the earth mover distance on the diffusion-based space to explore patient-to-patient variation. PhEMD lies in its dependence on diffusion maps and pseudo-time estimates, presupposing a continuous cellular progression in single-cell RNA sequencing data. This assumption may restrict its applicability in analyzing heterogeneous cell populations in whole organ samples, where such a linear cellular continuum may not exist. PILOT (Patient level distance with Optimal Transport) [19] investigates the sample-based analysis in multi-scale single-cell and pathomics data using optimal transport. However, it assumes that single-cell experiments have been previously pre-processed (clustered) and may be limited cell-type labels if unavailable.

In this study, we propose Quantized Optimal Transport (QOT), a fast and robust method of learning the distance matrix between samples from single-cell datasets using the OT-based Wasserstein distance. The schematic design of our proposed QOT method is shown in Figure 1. We demonstrate the promise of the proposed method through an empirical study on analyzing four different real world single-cell omics datasets.

**Figure 1:**
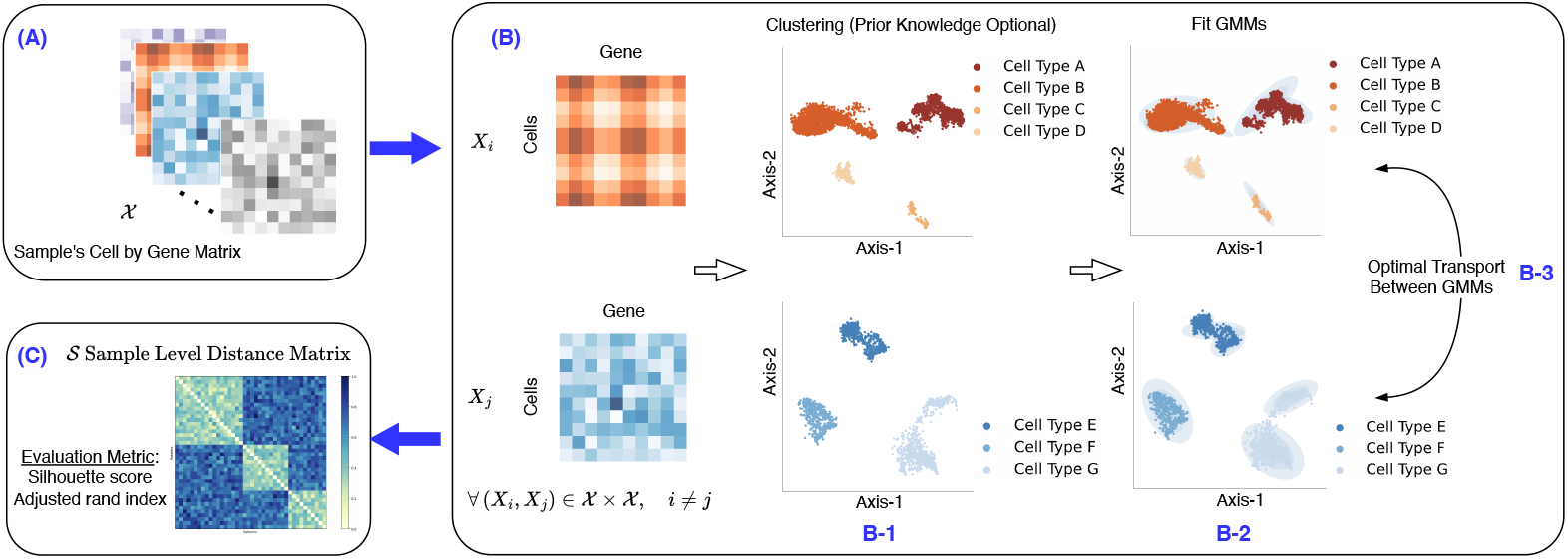
Schematic design of the proposed QOT algorithm. **(A)** Given the sample matrix, Space 𝒳, each matrix within it represents the single-cell gene expression for a sample. **(B)** The Quantized Optimal Transport (QOT) algorithm processes this data as follows: (B-1) First, it clusters each sample matrix based on its cell type information. If no prior knowledge is available, the clustering is performed using HDBSCAN. (B-2) Subsequently, a Gaussian Mixture Model (GMM) is used to model each cluster as a Gaussian distribution. (B-3) Finally, to derive the distance matrix between two samples, the OT-based Wasserstein distance is computed between their respective GMMs, where two versions of the calculation methods are presented. **(C)** The result of this computation yields the sample level distance matrix and is available for further downstream analysis.

### Our contributions can be summarized as follows

- **A New Method: QOT.** We propose Quantized Optimal Transport (QOT) to analyze single-cell gene expression data and quantify similarities and differences between samples. Our method can work with or without prior knowledge (cell type information), making it flexible and practical.
- **Empirical Performance Analysis.** We empirically evaluate QOT on four real-world datasets by comparing its performance against PILOT, which has the same setting in this problem formulation. Notably, our method achieves better accuracy than PILOT.
- **Complexity Analysis.** provide a detailed theoretical analysis of our proposed method’s time and memory complexity, offering insights into its efficiency.
- **Software.** The QOT code is available at https://github.com/PennShenLab/QOT.

The rest of the paper is organized as follows. In the **Material and Methods** Section, we first describe the real-world datasets utilized in this study. Following this, we review the background of several key components fundamental to our proposed method, ending with an in-depth description of the Quantized Optimal Transport. In the **Experiment Setup and Results** Section, we present the experiment setup and outcomes from applying QOT to real-world datasets, followed by the downstream trajectory analysis and theoretical analysis of computational complexity. The **Conclusion** Section summarizes our findings, discusses the limitations, and identifies possible areas for future research.

## 2 Material and Methods

### 2.1 Dataset

Table 1 summarizes the four datasets used in this study. Myocardial infarction is a leading cause of death worldwide. We used the Myocardial infarction scRNA-seq dataset generated from the study of [20]. It contains 115,517 cells from 20 samples, of which 13 are healthy controls and remote zones, and 7 are ischemic zones.

**Table 1:** Dataset information used in study.

|  | Myocardial Infarction | PDAC | Kidney IgAN (Tubule) | Kidney IgAN (Glomeruli) |
| --- | --- | --- | --- | --- |
| Type | scRNA-seq | scRNA-seq | Pathomics-seq | Pathomics-seq |
| Cells/Structures | 115, 517 | 57,530 | 64,493 | 24,227 |
| Samples | 20 | 35 | 634 | 634 |
| Cell_types | 11 | 10 | 15 | 14 |
| Classes | 2 | 2 | 3 | 3 |

Chronic kidney disease, a prevalent condition among adults, is a progressive illness with no known cure and is associated with significant morbidity and mortality. We used the pathomics data of kidney IgAN biopsies with two different morphological features, tubules and glomeruli, from [21]. The dataset is preprocessed using method form [4]. Both datasets are labeled based on their glomerular filtration rate (GFR): normal (GFR *>* 60), reduced (30*<*GFR*<*60), or low (GFR*<*30). Both datasets contain 400 normal samples, 177 reduced samples, and 57 low samples. The kidney IgAN (tubules) dataset has 64,493 structures and 15 cell types. The kidney IgAN (Glomeruli) contains 24,227 structures and 14 cell types.

Pancreatic ductal adenocarcinoma (PDAC) is the most common malignancy of the pancreas. We used the dataset from [22]. It has 11 healthy cohorts and 24 PDAC patients. The data is analyzed using Seurat and re-annotated cell clusters using the same marker genes by [19]. In total, it has 57,530 scRNA-seq cells from 10 cell types. Detailed information is shown in Table 1.

### 2.2 Methods Background

In this section, we briefly introduce the background knowledge required to understand our method. We first present Gaussian Mixture Model (GMM), which is used as a parametric tool to model our single cell omics data (**Figure 1, B-2**). After that, we present Optimal transport, which is used to define the distance between two samples (**Figure 1, B-3**).

#### 2.2.1 Gaussian Mixtxure Model

A Gaussian mixture model [23] is a weighted sum of M component Gaussian densities and it has the following form:

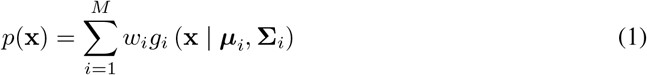

where

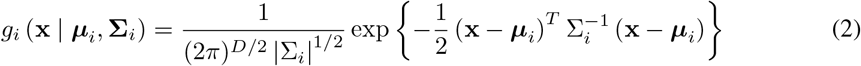

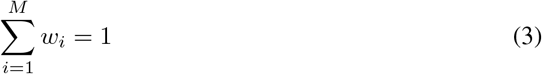

Here, *x* is a *N* -dimensional vector; *µ*_*i*_ and Σ_*i*_ is the mean and covariance of the *i* th guassian function; *w*_*i*_ is the coefficient of the mixture weights.

#### 2.2.2 Optimal transport

The Optimal Transport problem calculates the minimal cost required to transform one probability vector into another, serving as a measure of the similarity between the two vectors. For two multi-variate distributions, with position 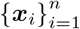 and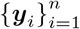. The discretized distributions are:

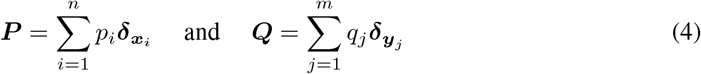

where *δ*_***x***_ denotes a Dirac delta function placed at a location ***x*** ∈ R^*n*^. The optimal transport problem could then be defined as:

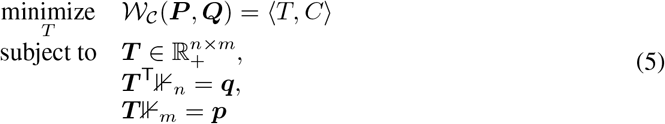

where the ***C***_*ij*_ = ∥***x***_*i*_ ***x***_*j*_∥^2^ is the cost matrix and ***T*** is the transportation plan we want to find that minimized the total transportation cost subjected to two marginal constrains.

To solve this problem, it suffers from the time complexitity 𝒪(*n*^3^ log *n*) and the curse of dimensionality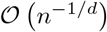. However, by introducing a regularized term, we could turn this problem into a strictly convex function with ideal statistical property. This sinkhorn distance [7] is defined as:

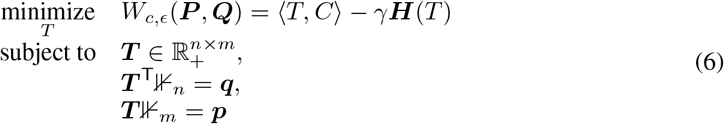

where ***H***(***T***) is the entropy of the transport plan matrix ***T*** and is given by 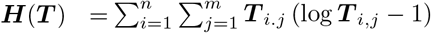. Moreover, to cancel the effect of the bias term, the sinkhorn divergence[24] is proposed.

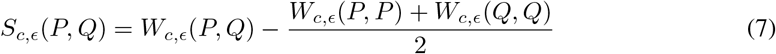

It has several advatanges: it could be interpresed as interpolating between the MMD and the Wasserstein Distance, it has complexity 𝒪 (*Ln*^2^)etc.

### 2.3 Proposed QOT Method

**Figure 1** shows the schematic design of our proposed Quantized Optimal Transport (QOT) method. The main problem is to obtain the sample level distance matrix from the cell by the gene expression matrix of each sample. Assuming there are *L* samples, yielding 𝒳 = {*X*_1_, *X*_2_ …, *X*_*L*_ }. *X* = ∈ [*x*_*ij*_] ∈ ℝ^*m×n*^ is the cell-by-gene expression matrix, where *m* is the number of cells and *n* is the number of genes, and *x*_*ij*_ represents the expression level of gene *j* in cell *i*. The task is to obtain the ∈ sample level distance matrix *S* = [*s*_*ij*_] ∈ ℝ^*L×L*^, where *s*_*ij*_ represents the distance between *i*^th^ and *j*^th^ samples.

To accurately obtain a sample level distance matrix from single-cell data, it is essential to meet two key criteria. First, an appropriate metric is needed that can address variations in cell counts across samples and the proportions of different cell types, which are often used as indicators of pathological changes or developmental differences. The Wasserstein distance is suitable in this context as it encompasses both aspects. Second, given the large single-cell datasets, the computational demand for calculating the Wasserstein distance should be carefully managed. To address this issue, our approach begins with quantization, which simplifies the complex data into a more structured, discrete format. Following this, our QOT Method transforms cell-by-gene expression matrices into parametric Gaussian Mixture Models (GMMs), facilitating efficient Wasserstein distance computations between samples.

Two primary steps are involved in the proposed Quantized Optimal Transport (QOT) Method for generating sample level distance matrix: 1. Acquisition of Gaussian mixture representations for each sample (**Figure 1, B-2**). 2. Computation of Wasserstein distances based on these Gaussian mixture representations (**Figure 1, B-3**). Within this framework, we differentiate between two QOT methodologies. The QOT-Exact method adopts a parametric approach, approximating the Wasserstein distance between two samples by restricting the set of possible coupling measures also to be Gaussian. In contrast, the QOT-Adaptive method is characterized by its focus on extracting key distributional features (i.e., emphasizing central tendencies). It computes the Wasserstein distances based on the centroids of the Gaussian mixtures, integrating metrics such as angular differences, spatial distances, and the alignment of covariances. Therefore, QOT-Exact aims to approximate the true Wasserstein distance between two samples, while QOT-Adaptives focuses more on the characteristics of each cell type.

#### 2.3.1 Gaussian Mixture Representation

In this section, we focus on clustering and characterizing each sample’s cell type using a Gaussian Mixture Model. We view samples as distributions of cells and cluster each sample by their cell type, leading to the formation of clusters 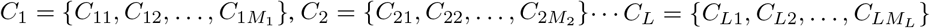. Each cluster’s empirical measure is determined by

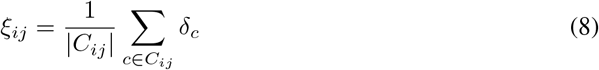

where *C*_*i*_ represents the cluster represetation of matrix *X*_*i*_, *C*_*ij*_ is defined as the *j*^th^ cluster of *i*^th^ sample, and *M*_*i*_ is the total number of cell type for *i*^th^ sample, *δ*_*c*_ is the Dirac measure centered at each data point *c*, |*C*_*ij*_| is the number of points in *C*_*ij*_.

Next, we apply a Gaussian Mixture Model (GMM) to each cluster *C*_*ij*_, The quantization operator *Q*(·) maps each cluster’s empirical measure to a Gaussian Mixture Model (GMM):

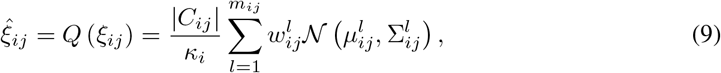

where 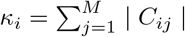 is the total number of poins in *i*-th sample; 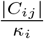 represents the proportion of *j*-th cluster in *i*-th sample; *m*_*ij*_ is the number of Gaussian components for cluster 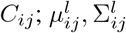 and 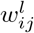, representing the mean, covariance, and weight of the *l*-th Gaussian component in *C*_*ij*_.

Next, a re-labeling step simplifies the component indexing. For each sample, each Gaussian components originally indexed by 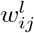, is re-labeled as *w*_*i,t*_, with *t* ranging from 1 to *N*_*i*_. *N*_*i*_ is the number of gaussian components of *i*-th sample:

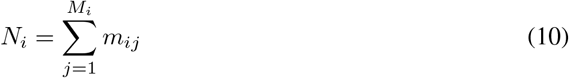

where *M*_*i*_ is the total number of clusters and *m*_*ij*_ denotes the number of Gaussian components in cluster *C*_*ij*_. Weights are then normalized to

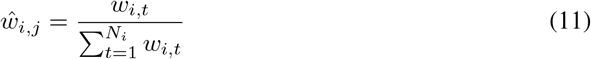

yielding a final model for each re-labeled, normalized cluster:

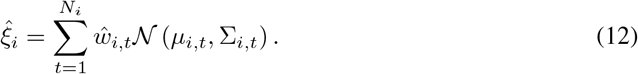

Here, 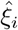 is the final Gaussian Mixture Representation of *i*-th sample; *µ*_*i,t*_ and Σ_*i,t*_ are the mean and covariance of the *t*-th Gaussian component of *i*-th sample. The details of the step by step algorithm are summarized in Algorithm 1.

#### 2.3.2 Quantized Optimal Transport-Exact

This section focuses on calculating the Wasserstein distance for two samples between their Gaussian Mixture Representation. Then, we could loop through all the samples to obtain the sample level distance matrix. To compute the Wasserstein distance between GMMs, a typical method is to use sampling strategy to discretized the GMMs and then solve the OT problem. However, the complexity of computing such an OT problem can increase based on the resolution of the discretization. In our approach, we compute the Wasserstein distance of two GMMs by restricting the set of possible coupling measures also to be Gaussian mixtures, i.e. we compute the optimal transport on the submanifold of Gaussian mixture distributions ([25]). The Wasserstein distance between quantized GMMs of two distinct samples is given by

##### Algorithm 1

Gaussian Mixture Representation for gene by gene expression matrix

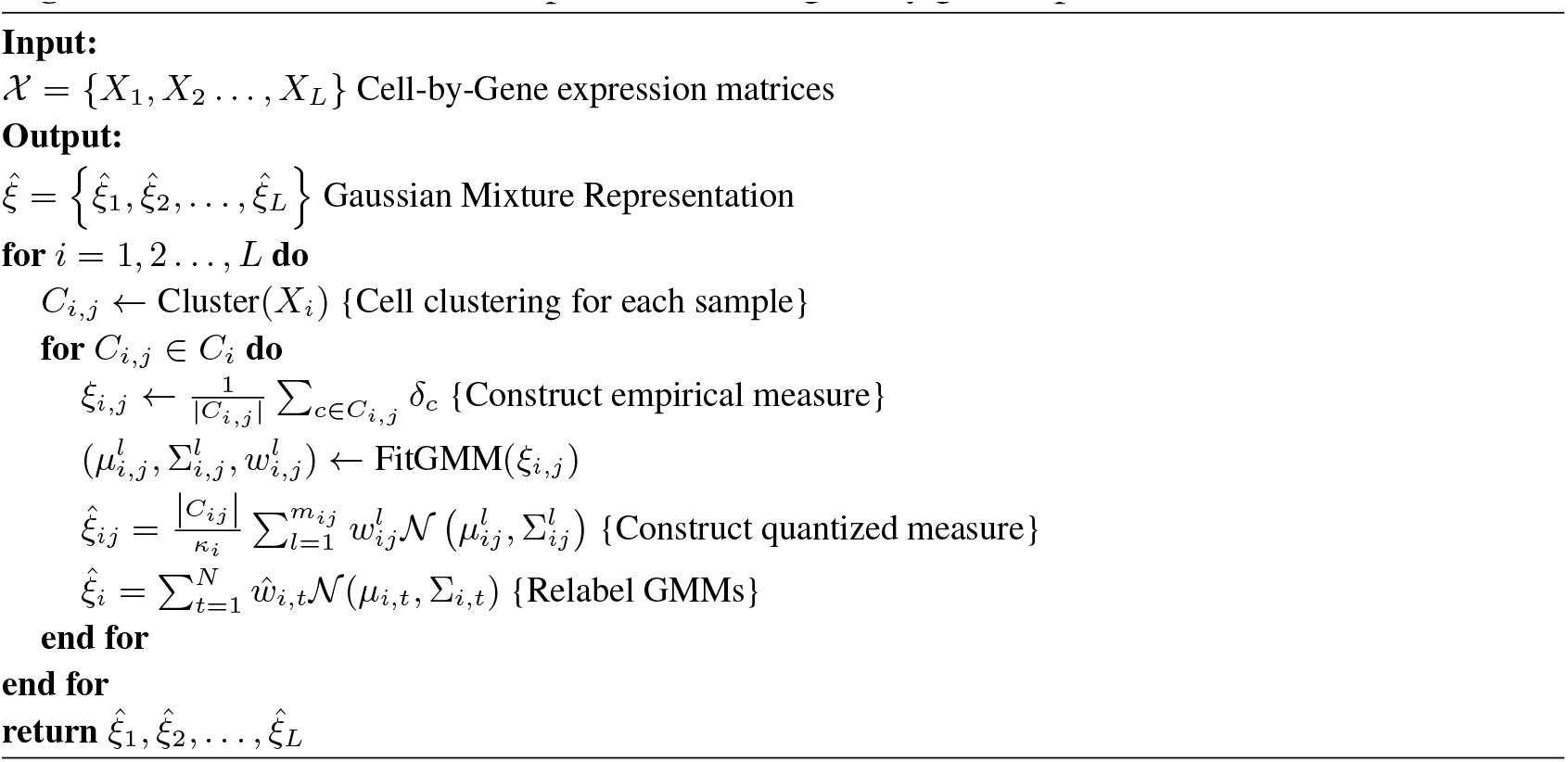

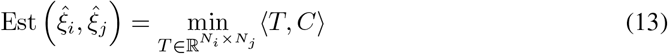

under the conditions

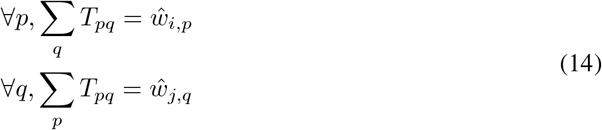

with the cost matrix defined by

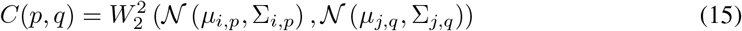

Here, *µ*_*i,p*_ and Σ_*i,p*_ denote the mean and covariance of the *p*-th Gaussian component of sample *i*, and similarly for sample *j*. Notice that the wasserstein distance between two gaussian distribution yields a closed form solution:

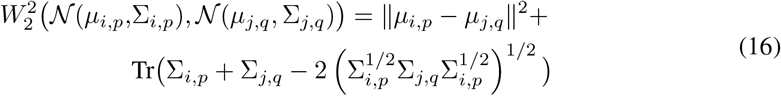

The details of the step by step algorithm are summarized in Algorithm 1.

#### 2.3.3 Quantized Optimal Transport-Adaptive

Here, we proposed an alternative way for the quantized optimal transport. Opposed to computing the parametric distance to approximate true Wasserstein distance from two samples, in this formulation, we calculate the cost matrix based on the euclidean or cosine similarity between GMM’s centroid along with the covariance:

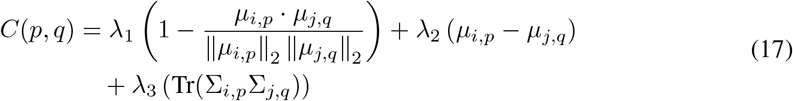

where the first term is the cosine similarity, the second term is the euclidean distance, and the third term quantifies the alignment of multidimensional variances, representing the geometric overlap of the distributions defined by two covariances. The details of the step by step algorithm are summarized in Algorithm 1.

##### Algorithm 2

Quantized Optimal Transport - Exact

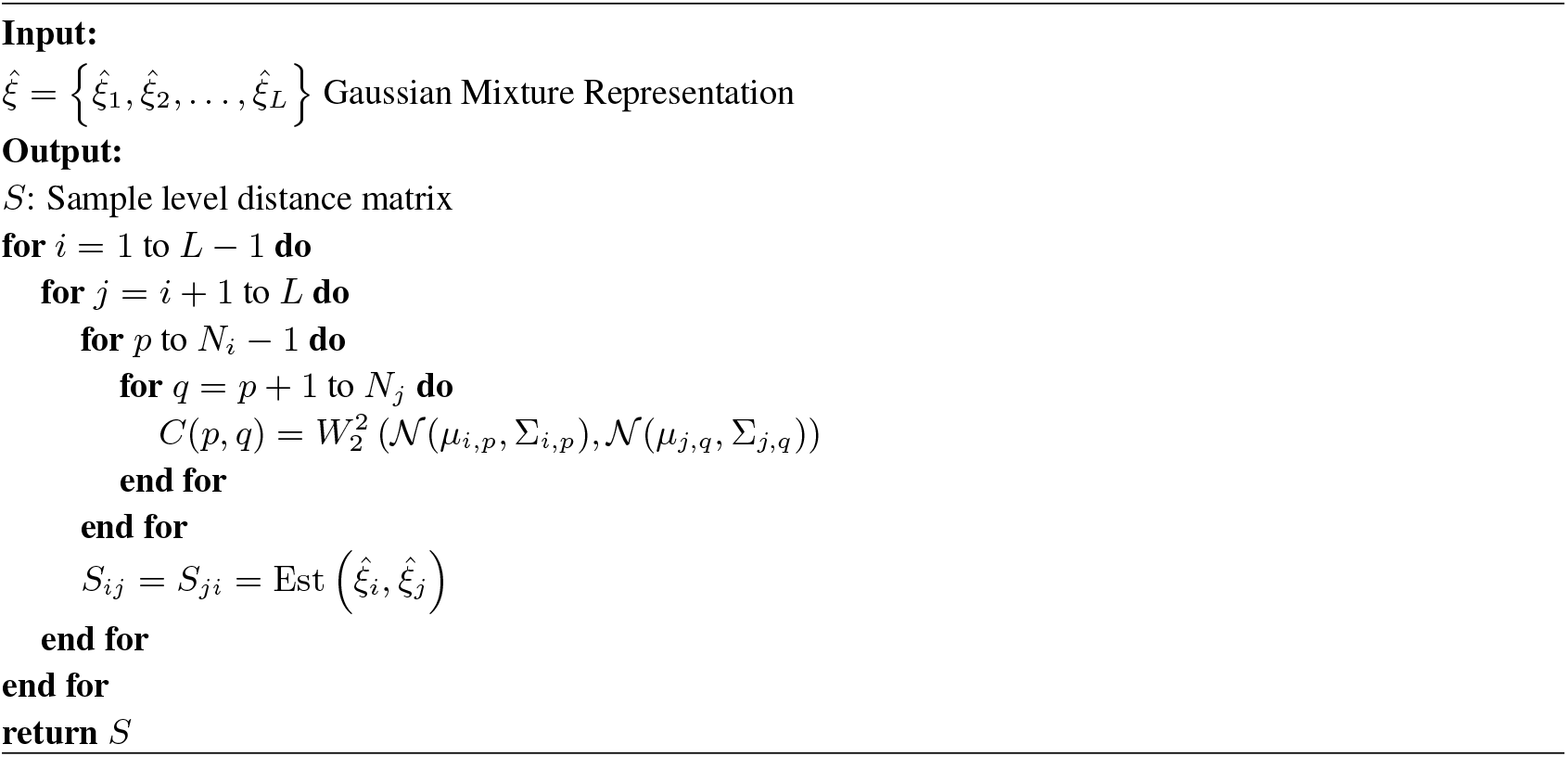

#### 2.3.4 Control the Quantization Error

In this section we discuss methods to control quantization error for better approximation. Choosing the proper clustering method is essential. If we have prior knowledge about the dataset, we can use cell type for clustering. Unsupervised clustering methods like centroid-based, graph-based, and density-based are options without this information. For single-cell data, we recommend using the density-based method HDBSCAN ([26]). This choice is due to the large number of cells in each sample and the importance of density in our data. The chosen quantization method could be based on specific tasks. In our methodology, after securing the clustering results, we applied the Gaussian mixture model (GMM) to capture each cluster’s nuances comprehensively. Determining the optimal number of Gaussian distributions for a fitting representation was crucial to this process. Intriguingly, given the nature of optimal transport in our approach, the specific number of Gaussians chosen to represent a cluster is independent of accuracy, making overfitting less of a concern. However, there is a trade-off to consider: using an excessive number of Gaussians might lead to computational inefficiencies without corresponding gains in precision. We employ the elbow method to determine the ideal number of Gaussian distributions for the GMM, which helps make an informed choice by plotting the explained variation against the number of clusters.

#### 2.3.5 Link to Relevant Prior Work

Recently, Patient level distance with Optimal Transport (PILOT) ([19]) has been proposed to investigate the patient level distance using the Wasserstein distance. Compared to PILOT, our method (QOT-Adaptive) could be seen as the generalized version of PILOT. In PILOT, the author used the centroid of each cell type to calculate the Wasserstein distance. It could be seen as QOT-Adaptive with Gaussian components to be 1 for all the clusters with cosine similarity only. Due to the nature of the dataset, it may not be sufficient to only consider the angle between the clusters. Therefore, our method considers both magnitude, angle, and cluster shape. Moreover, PILOT requires the true cluster (cell type information) that will limit its usage in another dataset. This is because the cost matrix of the PILOT is based on the pair-wise cell type. In another way, our method only used the cell-type information to compute the Gaussian mixture model. The cost matrix is defined w.r.t the Gaussian component. For example, if two samples have ten cell types and a 3-component Gaussian mixture model fits each cell type. The cost matrix for the PILOT will be a ten by ten matrix and thirty by thirty for the QOT method. When the cell-type information is unavailable, we recommend the user perform unsupervised clustering of the data via HDBSCAN and then fit the Gaussian mixture model from the inferred cluster.

##### Algorithm 3

Quantized Optimal Transport - Adaptive

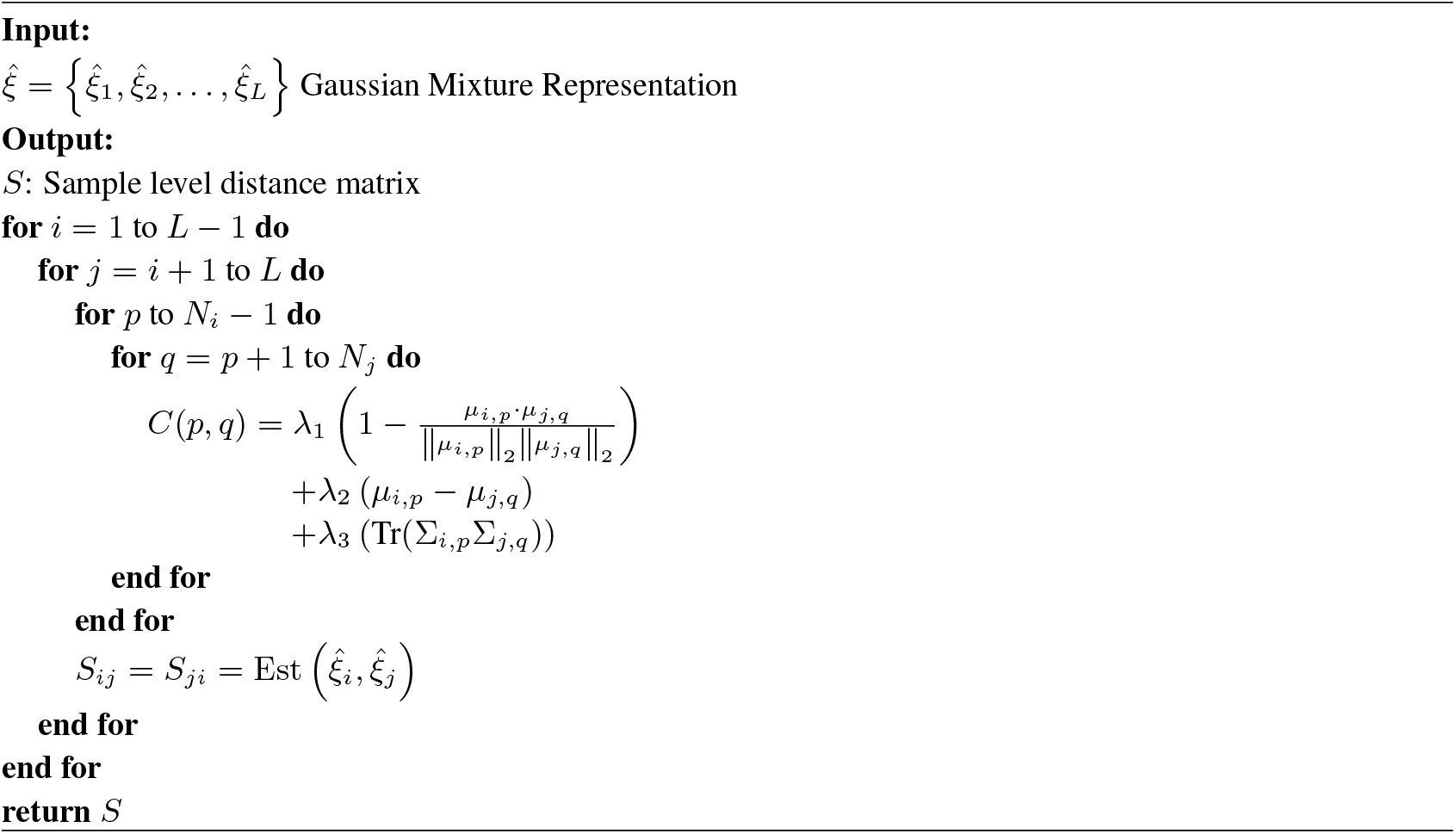

## 3 Experiment Setup and Results

### 3.1 Baseline and Evaluation Metric

We empirically evaluate our method on the same datasets used in PILOT [19], which has demonstrated desirable results. In this work, we benchmark against the PILOT and showcase the enhanced capability of our method.

The Silhouette score evaluates the sample level distance matrix (i.e., how well the distance matrix aligns with their true labels). It is defined as:

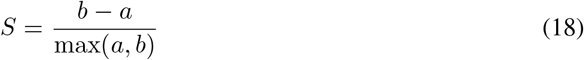

where *a* is the mean distance between a sample and all other points in the same cluster, and *b* is the mean distance between a sample and all other points in the nearest cluster that the sample is not a part of. This score ranges from -1 to 1, where a high value indicates that the sample is well-matched to its cluster and poorly matched to neighboring clusters. In the experiment part, as shown in Table 2, we provide two versions of Silhouette. As the PILOT and QOT methods both output the distance matrix, the first index is named Sil. Directly use the inferred distance matrix to calculate the silhouette score. The second is named Sil.Pilot, the PILOT version of the silhouette score. It treated the inferred distance matrix as the vector representation of each Sample and used the cosine similarity of the inferred distance matrix as the input of the silhouette score.

**Table 2:**
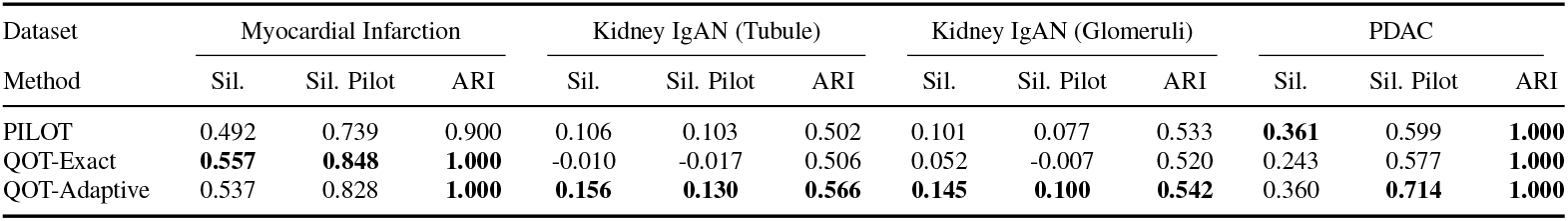
Performance Comparison across different datasets. QOT method ranks best for Myocardial Infarction, Kidney IgAN (Tubule), and Kidney IgAN (Glomeruli). Notice in the PDAC dataset, we fit a single Gaussian distribution to each cell type, and therefore, the result should be almost the same as the PILOT method under Sil. (PILOT 0.361, QOT-Adaptive 0.360).

The adjusted rand index (ARI) is used to evaluate how well different methods maintain connections between samples linked to their labels; we performed unsupervised clustering on a k-nearest neighbor (kNN) graph created among the samples, utilizing the Leiden algorithm for this process. It is defined as:

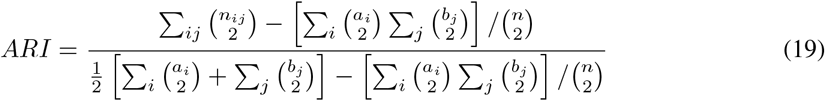

where,*n*_*ij*_ is the number of objects in both the *i*-th group of the first clustering and the *j*-th group of the second clustering, *a*_*i*_ is the number of objects in the *i*-th group of the first clustering, *b*_*j*_ is the number of objects in the *j*-th group of the second clustering, *n* is the total number of objects, 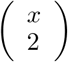 represents a binomial coefficient, calculating the number of unique pairs that can be formed from *x* objects. The algorithm is implemented in Python and runs on a system with an x86_64 architecture, Intel(R) Xeon(R) CPU operating at 2.20GHz, and 51GB of RAM. QOT is based on the emd2 function from the POT library ([27]). The computation of Silhouette and adjusted rand index (ARI) are based on sklearn.metrics.silhouette_score and sing the sklearn.metrics.adjusted_rand_index from scikit-learn v1.2.2.

### 3.2 Evalutation of Sample Level Distance Matrix

We first evaluate QOT-Exact and QOT-Adaptive on four public datasets with known sample status labels. We evaluate our method based on the silhouette score and the adjusted rand index. As shown in Table 2, for the Myocardial Infarction dataset, a four components Gaussian mixture model fits each of the cell-type, QOT-Exact ranked first under Sil index, and QOT-Adaptive ranked first under Sil.Pilot and ARI. For both Kidney IgAN datasets, each cell type is fitted by a three-component Gaussian mixture model. On the Kidney IgAN (Tubule) dataset, QOT-Adaptive ranks first under Sil., Sil.Pilot, and ARI, while QOT-Exact rank last under Sil., Sil.Pilot and second in the ARI. On the Kidney IgAN (Glomeruli) dataset, QOT-Adaptive ranks first under Sil., Sil.Pilot, and ARI. The PILOT ranks second under Sil., Sil.Pilot, and ARI. QOT-Exact is the last under Sil. Sil.Pilot, and ARI. Finally, on the PDAC dataset, each cell type is fitted by a one-component Gaussian mixture model. In the PDAC dataset, QOT-Adaptive ranks first in the Sil.PILOT, PILOT ranked first in the Sil. Both three methods tied up in 1 under the ARI index. Distance Matrix with best Sil. among QOT-Exact and QOT-Alternative are shown in Figure 2 and Figure 3. In the PDAC dataset, PILOT and QOT-Adaptive use each cell type’s mean to calculate the Wasserstein distance. Therefore, when comparing the output distance matrix directly (Sil.), there should be a slight difference due to the difference in how to choose the mean within cell type (In PILOT, the author directly calculates the mean, and in QOT-Adaptive, we fit the parametric model and then extract the mean) This is confirmed by the fact under Sil. Index, PLIOT achieves 0.361, and QOT-Adaptive achieves 0.360. However, the difference may be magnified when we do not directly compare the distance matrix (i.e., view the matrix as the vector representation).

**Figure 2:**
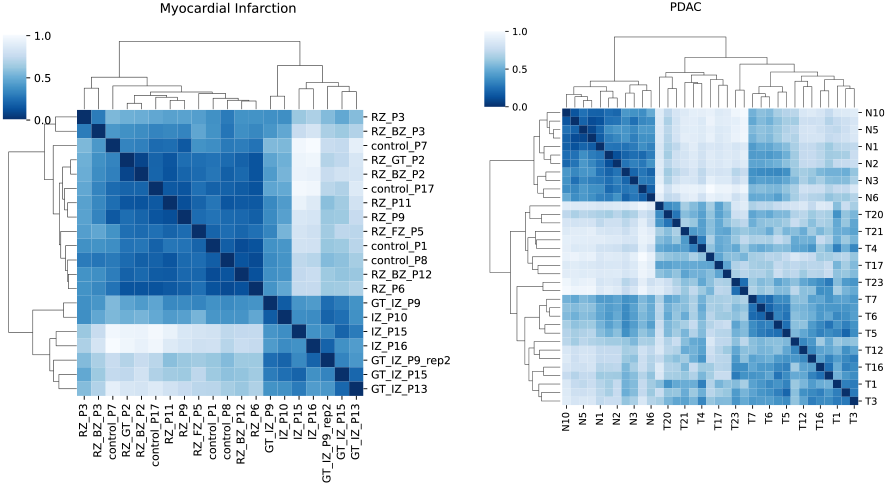
Distance Matrix with best Sil. among QOT-Exact and QOT-Alternative for the rna-sequence dataset. Left: Myocardial Infarction dataset; Right: PDAC dataset. Legend represents the distance between two samples.

**Figure 3:**
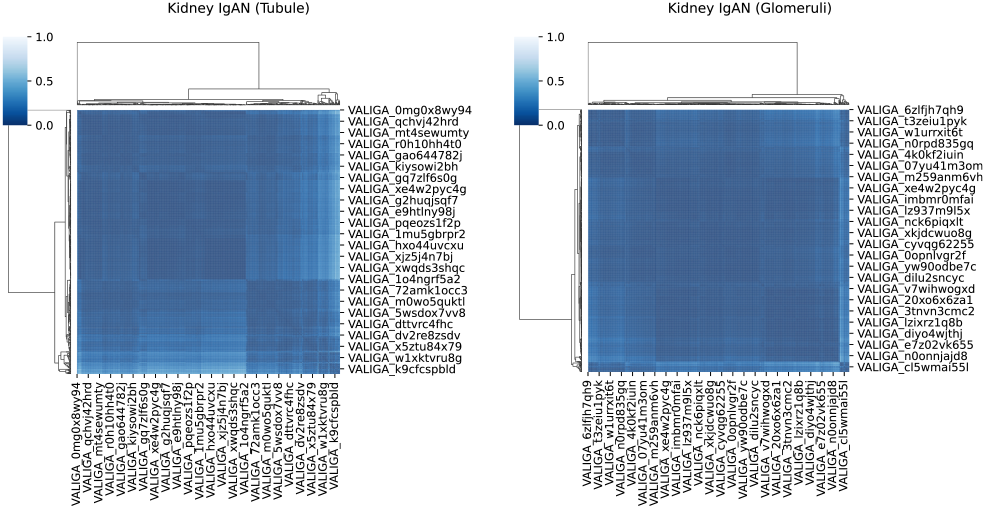
Distance Matrix with best Sil. among QOT-Exact and QOT-Alternative for the pathomic dataset. Left: Kidney IgAN (Tubule) dataset; Right: Kidney IgAN (Glomeruli) dataset. Legend represents the distance between two samples.

Also, when the number of Gaussian components for each cell type is greater than one, we argue that we obtain a more detailed description in QOT-Adaptive than in PILOT since each cell type is delineated more than its mean. This fact is further confirmed in the other three datasets: Myocardial Infarction, Kidney IgAN (Tubule), and Kidney IgAN (Glomeruli), where the performance is improved.

### 3.3 Sample Level Trajectory Analysis

One downstream analysis for a sample level distance matrix is to uncover the progression of a disease. This is achieved by embedding samples into a two-dimensional space using PHATE (Potential of Heat-diffusion for Affinity-based Trajectory Embedding, as described by [28]). The results reveal a strong correlation between the progression of the disease and the pattern observed along the curve in the embedding space. This implies that the sample level distance matrix could effectively discern the essential phenotypic distinctions among samples. This downstream analysis method is also used in PILOT. Our method also shows improvement in this Trajectory task. We calculate the Area Under the Curve of the Precision-Recall curve (AUCPR) to quantify whether the generated trajectory follows the disease progression order. For datasets with two classes (Myocardial Infarction and PDAC), we use label 0 for a normal sample and 1 for the disease sample. For the dataset with three classes (Kidney IgAN (Tuble) and Kidney IgAN (Glomeruli), we use 0 for the normal sample, 1 for the reduced sample, and 2 for the low sample. Moreover, for both Kidney datasets, AUCPR is computed by treating each class as a separate binary problem (One-vs-Rest), calculating AUCPR for each, and then averaging these scores to get the overall performance. We compared the trajectory map using distance generated by PILOT and QOT methods, where the choice among QOT-Exact and QOT-Labeled is based on Sil. For Myocardial Infarction, we use QOT-Exact. For PDAC, Kidney IgAN (Tubule), and Kidney IgAN (Glomeruli), we use the QOT-Alternative. The results of the trajectory map are shown in Figure 4, and the AUCPR is shown in Table 3.

**Table 3:** Evaluation pf trajectory map across different datasets Using AUCPR.

|  | AUCPR |  |  |  |
| --- | --- | --- | --- | --- |
|  | Myocardial Infarction | Kidney IgAN (Tubule) | Kidney IgAN (Glomeruli) | PDAC |
| PILOT | 0.915 | 0.540 | 0.529 | **1.000** |
| QOT | **1.000** | **0.560** | **0.540** | **1.000** |

**Figure 4:**
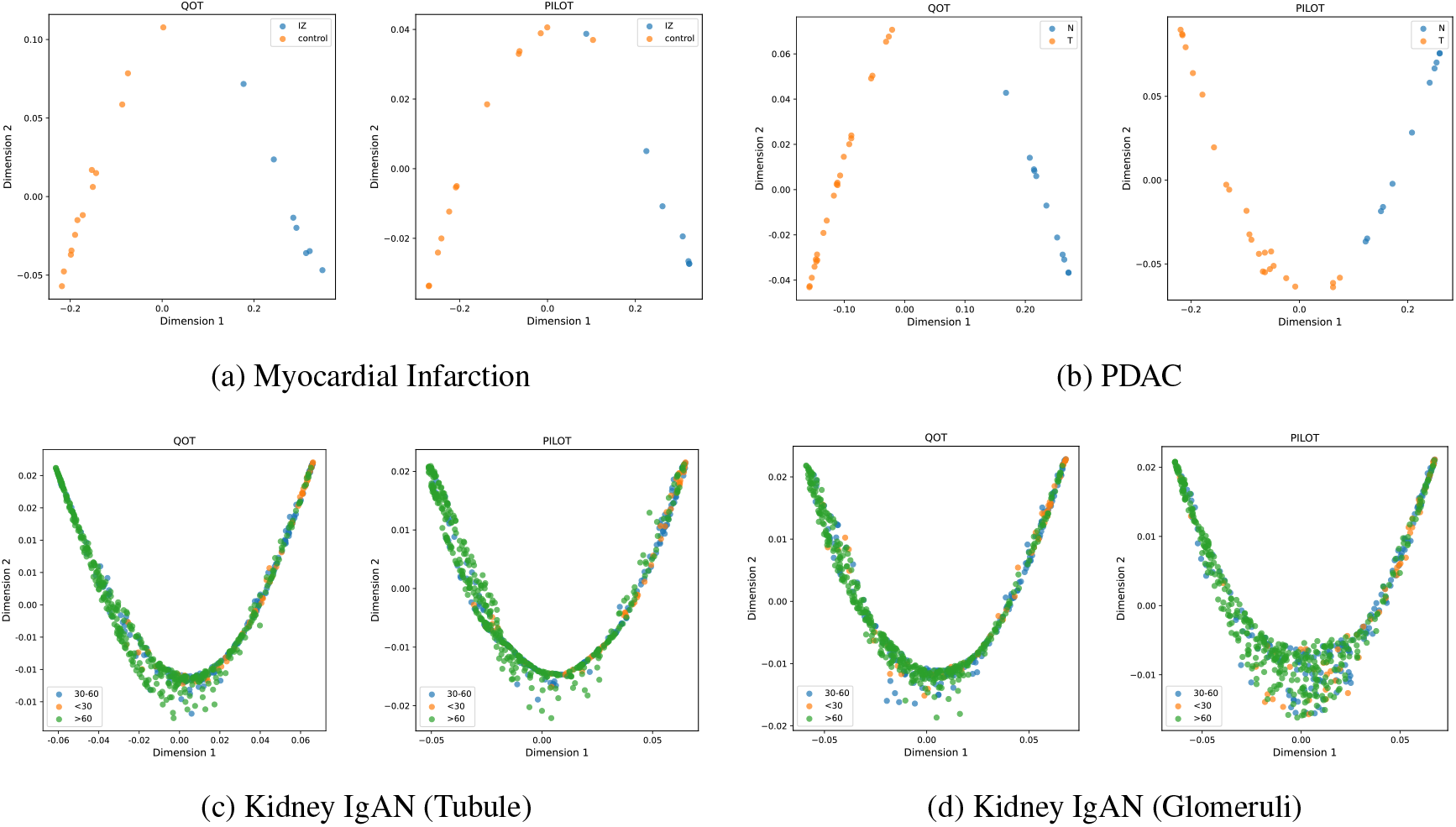
Trajectory Analysis for singe-cell and pathomic dataset. (a)-(d) are Myocardial Infarction dataset, PDAC dataset, Kidney IgAN (Tubule) dataset, and Kidney IgAN (Glomeruli) dataset. In each subfigure, left is the trajectory generated from QOT method distance matrix and right is the trajectory generated from PILOT method distance matrix.

The QOT method has the highest AUCPR score of 1 for the Myocardial Infarction dataset. Notice that, in Figure 4 (b), the PILOT method incorrectly placed an IZ (disease) sample between two control samples. In contrast, the QOT method has the correct disease progression order (normal then disease). For the PDAC dataset, the QOT and PILOT methods have the correct disease progression order (normal then disease). For the Kidney IgAN (Tubule) and Kidney IgAN (Glomeruli) datasets, the QOT method has a higher AUCPR score, yielding more accurate disease progression.

### 3.4 Complexity Analysis

To provide an accurate picture of the actual cost of the algorithm, we provide the time and space complexities of all the methods used here. Both complexities are evaluated based on the cost of computing between a pair of samples. Given two samples, each with cells of *N*_1_ and *N*_2_, we define *N* = max(*N*_1_, *N*_2_), both containing *D* features, *S* cell-type, and *K* to be the total number of Gaussian components for each sample. The results are shown in Table 4.

**Table 4:**
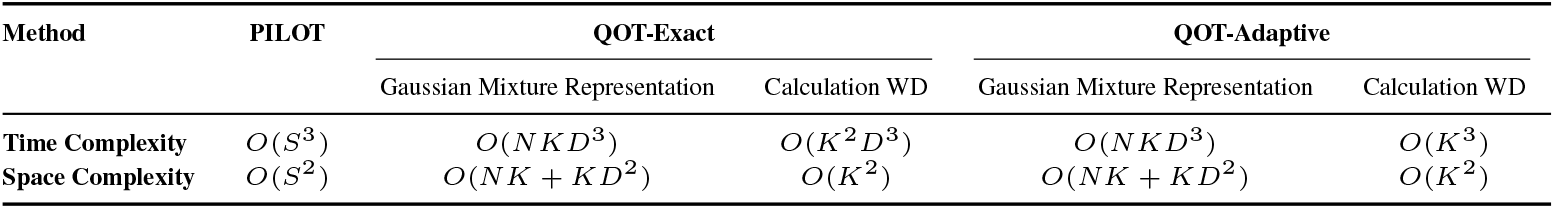
Comparison of time and space complexities between PILOT and QOT. Given two samples, each with cells of *N*_1_ and *N*_2_, we define *N* = max(*N*_1_, *N*_2_), both containing *D* features, *S* cell-type, and *K* to be the number of Gaussian components for each cell type.

| Method | PILOT | QOT-Exact |  | QOT-Adaptive |  |
| --- | --- | --- | --- | --- | --- |
|  |  | Gaussian Mixture Representation | Calculation WD | Gaussian Mixture Representation | Calculation WD |
| Time Complexity | $O(S^3)$ | $O(NKD^3)$ | $O(K^2D^3)$ | $O(NKD^3)$ | $O(K^3)$ |
| Space Complexity | $O(S^2)$ | $O(NK + KD^2)$ | $O(K^2)$ | $O(NK + KD^2)$ | $O(K^2)$ |

For the PILOT method, the time complexity is *O*(*S*^3^), and the space complexity is *O*(*S*^2^) because the Wasserstein distance is computed between the centroid of each cell type.

For the QOT-Exact method, the time complexity is *O* (*NKD*^3^ + *K*^2^ *D*^3^), where *O* (*NKD*^3^) comes from the GMM step, and *O* (*K*^2^*D*^3^) is for computing the Wasserstein Distance between two GMMs.

The space complexity aggregates to *O* (*NK* + *KD*^2^ + *K*^2^), where *O* (*NK* + *KD*^2^) comes from the GMM step, accounting for the storage of posterior probability and covariance matrix, respectively, and *O* (*K*^2^) is the space to store the cost matrix.

For the QOT-Adaptive method, the time complexity is *O*(*NKD*^3^ + *K*^3^) *O*(*NKD*^3^), where ^3^ comes from the GMM step, and *O* (*K*^3^)is for computing the Wasserstein Distance between two GMMs’s mean. The space complexity aggregates to *O* (*NK* + *KD*^2^ + *K*^2^), where *O* (*NK* + *KD*^2^)comes from the GMM step, accounting for the storage of posterior probability and covariance matrix, respectively, and *O* (*K*^2^)is the space to store the cost matrix.

In reality, we assume that *D* ≪*N* and *K*≪ *N*. Therefore, the Gaussian Mixture representation is linear concerning *N* in both time and space complexities. In computing the QOT, the QOT-Adaptive is superior to the QOT-Exact. Also, the total number of Gaussian components *K* is multiple of the cell types *S* since we fit the Gaussian mixture model for each cell type. Therefore, the actual time complexity of QOT-Adpative is a multiple of the PILOT. It aligns with the intuition that QOT-Adaptive and PILOT use the centroid-based method to calculate the Wasserstein distance. At the same time, QOT-Adaptive is more detailed in obtaining the centroid representation of each cell type.

## Conclusion

This study introduced an efficient method for handling large-scale single-cell data. We successfully applied this method to four real world single-cell and pathomics datasets, achieving accurate results while maintaining computational efficiency. Our method offers a distinct advantage over optimal transport based method, PILOT, which primarily rely on cell type information. By incorporating geometric perspective alongside percentage information, we can better differentiate between various diagnostic groups.

Our method is fundamentally rooted in the idea that Wasserstein distance can effectively capture data distribution, and the quantization step plays a pivotal role in reducing computational complexity while retaining crucial geometric information within the data. The complexity analysis demonstrates that our method has linear time and space complexities with respect to the sample size. We plan to refine this further by reducing the algorithm’s sensitivity to the number of dimensions, thereby improving its robustness.

We acknowledge that a straightforward application of fitting Gaussian Mixture Models (GMMs) directly to the data may not always yield optimal results, primarily due to the challenge of selecting the appropriate number of components. Our quantization step based on prior or data driven clustering offers an efficient strategy for uncovering inherent sample characteristics.

After obtaining the Gaussian Mixture Representation of the data, we provide two version of QOT method that focusing on different aspect of the data. QOT-Exact, which computed the Wasserstein distance constraint on Gaussian coupling offers a fast tool for characterizing these features, could be considered as the efficient method for approximating the wasserstein distance between two dataset. QOT-Adaptive, in the other hand, shifts the focus in Gaussian Mixture Model analysis to the centroids of Gaussian components, emphasizing central tendencies. It combines euclidean and cosine measures with covariance alignment, offering another perspective on the geometric relationships and orientations within the data.

The following are a few interesting future directions. Firstly, while our current methodology involves fitting a Gaussian Mixture Model (GMM) to each cluster within each sample, further investigation into more efficient and accurate methods of parametrizing the data could be considered. Additionally, the quantization step itself could potentially be directly incorporated into the computation of the Wasserstein distance. By distilling the given input data and compressing the samples, we could improve the efficiency of the Optimal Transport (OT) plan from a different perspective of our current algorithm.

## 4 Competing interests

No competing interest is declared.

## 5 Author contributions statement

Conceptualization, Z.W., Q.Z., J.C., and L.S.; Methodology, Z.W., Q.Z., and L.S.; Resources, P.O., J.W., and L.S.; Formal analysis, Z.W., Q.Z., S.Y., J.C., and L.S.; Writing-Original Draft, Z.W. and J.C.; Funding acquisition, P.O., J.W., and L.S.; Writing-Review and Editing, Z.W., Q.Z., S.Y., S.M., J.C., S.G., P.O., J.W., and L.S.

## 6 Acknowledgments

This work was supported in part by NIH grants U01 AG066833, R01 AG071470, and U19 AG074879.

## Notes

https://github.com/PennShenLab/QOT

### Competing Interest Statement

The authors have declared no competing interest.

